# NANOME: A Nextflow pipeline for haplotype-aware allele-specific consensus DNA methylation detection by nanopore long-read sequencing

**DOI:** 10.1101/2025.06.29.662079

**Authors:** Yang Liu, Hash Brown Taha, Qiuyang Zhang, Ziwei Pan, Christina Chatzipantsiou, Emma Wade, Thatcher Slocum, Lasya Karuturi, Yue Zhao, Shilpita Karmakar, Sheng Li

## Abstract

Nanopore long-read sequencing has expanded the capacity of long-range, single-base, and single-molecule DNA-methylation (DNAme) detection and haplotype-aware allele-specific epigenetic phasing. Previously, we benchmarked and ranked the robustness of seven computational tools for DNAme detection using nanopore sequencing. The top performers were Megalodon, Nanopolish, DeepSignal and Guppy. However, these algorithms exhibit lower performance at regions with discordant non-singleton DNAme patterns compared to genome-wide regions. Furthermore, long-read sequencing analysis of mammalian genomes requires higher computational resources than next-generation sequencing. To address these issues, we developed a NANOpore Methylation (NANOME) a consensus DNAme predictive model using XGBoost, which integrates the output of Megalodon, Nanopolish, and Deepsignal for analyzing data obtained using Oxford Nanopore Technologies (ONT). NANOME enhanced DNAme detection precision (mean square error) at single-base resolution by 11% and improved accuracy (F1-score) at single-molecule resolution by 2.4% for human B-lymphocyte European cell lines (NA12878). The consensus model also detected ∼200,000 more CpGs than all three tools. Combing variant calling and long-read phasing, NANOME can detect haplotype-aware allele-specific DNAme in known imprinting controls in resolved and previously unresolved regions. We conducted haplotype-aware methylation detection on the T2T genome for dataset NA12878, revealing significant variations in differentially methylated region (DMR) density between gap and non-gap regions. Overall, NANOME represents a significant step forward in DNAme detection and long-range epigenetic phasing, offering a robust and accessible tool for researchers studying the epigenome.

## Main

DNA methylation controls the development and differentiation of cells by influencing which genes are expressed or repressed [1]. Detecting changes in DNA methylation patterns can provide valuable insights into disease mechanisms and potential therapeutic targets [2, 3]. Nanopore sequencing is a third-generation sequencing technology that utilizes nanopores to sequence DNA strands [4, 5]. Unlike second-generation sequencing technologies, which require amplification of the DNA molecule, Nanopore sequencing directly reads the DNA molecule by threading it through a protein nanopore, which is integrated into an electrically resistant membrane. As the DNA passes through the nanopore, it causes a characteristic change in the electrical signal, which is used to identify the nucleotides in the DNA sequence [6]. This technology has several advantages over other sequencing technologies, including the ability to sequence long-reads of DNA without the need for amplification, real-time monitoring of DNA sequencing, and the potential to sequence a wide range of nucleic acid samples, including RNA, DNA, and modified bases. Nanopore sequencing has been used in a variety of applications, including microbial genome sequencing, cancer research, and metagenomics, making it a powerful tool for advancing genomic research [7–9].

Various analytical tools for DNA methylation detection from long-read nanopore sequencing data, such as Oxford Nanopore Technologies (ONT), have been recently developed and evaluated [10–18]. However, these algorithms have strict parameters, are available through different Nanopore Community software packages and dependencies, and pose entry barriers for developers and users. Further, each of these tools involves different preprocessing steps. For instance, DeepSignal [11] and Tombo [12] require a re-squiggle algorithm, Nanopolish [17] requires an index step following minimap2 alignment, and Guppy requires a post-processing module, such as fast5mod or gcf52ref to extract the site level outputs, and does not support read-level outputs [14]. Such issues can hamper the comparison of method performance, resulting in poor reproducibility. Another key experimental challenge is the large amount of input DNA required for ONT sequencing. Megalodon and DeepSignal have shown increased prediction accuracy and CpG completeness with higher read coverage, with the caveat of requiring more computational resources and longer running times for tera-base scale input data [7]. Recently, some improved model of methods proposed [19–25]. Improving the performance of algorithms developed to characterize epigenomic events for base modifications at the single-molecule level can significantly advance single-molecule omics studies [26, 27].

Although the reproducibility and replicability of computational pipelines are critical, they are often overlooked in research [28] since they are challenging to achieve. To address this issue, domain-specific languages (DSLs), such as Nextflow and Snakemake, which require programming knowledge, have been recently developed specifically for streamlining bioinformatics tasks [29, 30]. Recently, these DSL-based pipelines have been successfully applied to long-read sequencing [7]. Furthermore, nanoseq, a Nextflow-based pipeline, has been developed for benchmarking RNA transcript level analysis [31]. There are several cloud-based pipelines currently available, such as Katuali [32], NanoSPC [33], and nanopore-gpu [34], which use Snakemake, Nextflow, and Biodepot-workflow-builder (Bwb) to perform base calling and alignment. Nanopype [35] is an easy-to-use Snakemake-based pipeline that provides DNA methylation-calling for ONT nanopore sequencing data by integrating Nanopolish. However, Nanopype does not support graphics processing unit (GPU) software versions in Singularity and Docker [36] containers and can only be run on local or cluster HPC. METEORE [37] is a consensus method that uses a Snakemake-based pipeline to make methylation-calling predictions from ONT nanopore data. This method combines two or more tools, including Nanopolish, Tombo, DeepSignal, Guppy, Megalodon, and DeepMod. Still, it fails to integrate Megalodon and Guppy to pipeline, which have been proven to be top performers [38, 39], and requires prior basecalling of the input data. Take the advantage of haplotype resolved variant calling [40], PRINCESS [41], NanoMethPhase [42] and MethPhaser [43] are proposed to phases 5-methylcytosine from nanopore sequencing data and performs single nucleotide variant (SNV) calling, Structure Variant (SV) and phasing [44, 45].

Nanopype and METEORE also only support haplotype-aware methylation solution to Nanopolish methylation call results, and the phased methylation performance of PRINCESS has not been evaluated on real word datasets [46]. EPI2ME Labs (https://labs.epi2me.io/wfindex) is an intuitive, multi-platform desktop environment developed by ONT that supports and develops open-source pipelines for nanopore sequencing data analysis. They have recently developed an all-in-one human variation Nextflow workflow, wf-human-variation (https://github.com/epi2me-labs/wf-human-variation). Still, this workflow only supports ONT-developed methylation tool outputs, and there is no Continuous Integration and Continuous Delivery (CI/CD) automation test features for Nextflow pipeline.

Recent studies have shown that no single methylation-calling tool can achieve the best performance across all metrics [37, 38, 47], making the development of a pipeline that supports DNA methylation-calling tools highly desirable. The key challenge faced by the end-to-end DNA methylation pipeline is its heterogeneous methylation-calling algorithms, which can be based on statistical, machine learning, deep learning, or consensus model algorithms. Additionally, these tools require varying hardware resources, with some needing GPU for acceleration (Guppy, Megalodon, DeepSignal, and DeepMod) while others require more CPUs and larger memory (DeepSignal and DeepMod). There has been little or no discussion about cloud computing support. Moreover, recent whole-genome performance reports show that no single tool is satisfactory across all metrics [38, 42].

To address this, we have developed the first DNA methylation-calling pipeline, termed NANOpore Methylation (NANOME), which integrates the state-of-the-art best performers and provides a consensus result that covers more CpGs than any single algorithm. Our pipeline is a reproducible, end-to-end solution for whole-genome DNA methylation-calling for nanopore sequencing data. Furthermore, the proposed pipeline is robust and user-friendly and includes cloud storage, computing, and real-time analysis (**Figure 1**). By accessing NANOME pipeline, users can run a single command line to analyze whole-genome high-throughput sequencing data.

**Figure 1.**
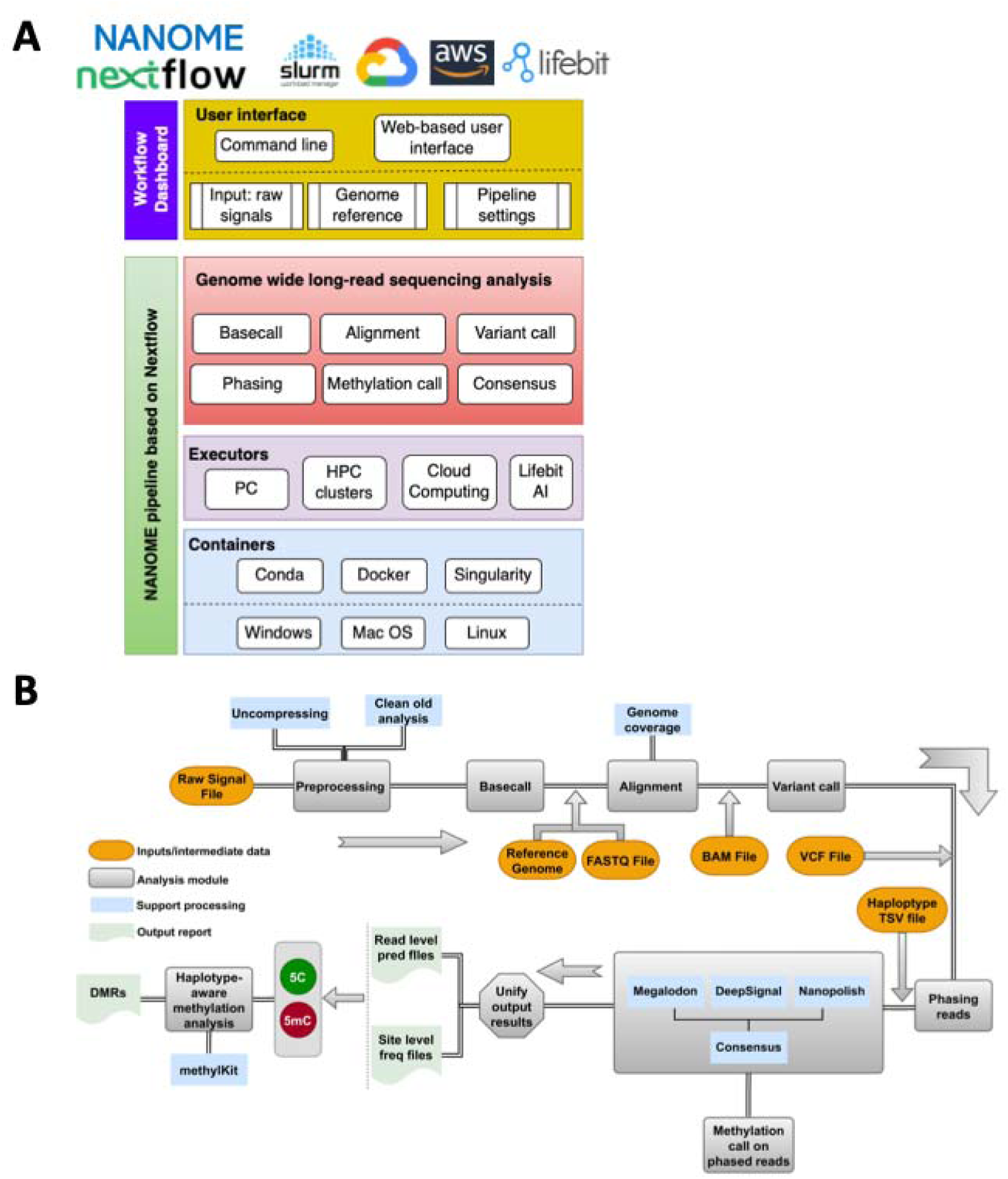
Architecture and workflow of the NANOME pipeline designed for DNAme detection on Oxford Nanopore long-read data. (**A**) Architecture of cloud compatible pipeline. (**B**) Workflow of NANOME pipeline. The pipeline offers flexibility to the user who choose to run a different DNAme tools after the preprocessing stage.

## Results

### NANOME:a fully automated pipeline for base- and methylation-calling on nanopore long-read data

We designed NANOME as a DNA methylation-calling pipeline to streamline long-read nanopore DNA sequencing data processing and analysis (**Figure 1A-B**). Our pipeline uses nanopore sequencing raw signal data in fast5 format and a reference genome as inputs. NANOME consists of several modules, including pre-processing, basecalling, alignment, methylation-calling, and generation of a consensus model. It integrates state-of-the-art software packages: Megalodon [10], Nanopolish [17], DeepSignal [11], Guppy [18], Tombo [12], DeepMod [15], and consensus model NANOME. Different tools require distinct alignment steps to align the Guppy basecalled results to a reference genome. Methylation-calling modules in Guppy, Megalodon, DeepMod, and Nanopolish already include minimap2 in their methylation calling module or execute the minimap step before methylation calling, while DeepSignal and Tombo use re-squiggle [11]. This design ensures consistency in processing nanopore data, enabling the capability of consensus of different computational methods.

We tested dynamically on three tera scale size datasets, NA12878, NA19240 and NA24385 (in **Methods**). NANOME features with state-of-the-art pipeline comparisons are listed in **Table 1**. Specifically, NANOME allows data processing through containers, including Conda, Docker, and Singularity on different operating systems such as Windows, Mac OS, and Linux, and can be executed on the cloud; all tools were containerized to address language-specific package dependencies. Furthermore, NANOME can run multiple tools in above in parallel, facilitating the direct comparison of their performance, which is particularly useful for evaluating and applying methylation-calling tools. The deep learning model adjustment for DeepSignal, DeepMod, and Megalodon can be conducted in the command line or web interface, configured by running the model on HPC clusters or the cloud.

**Table 1.**
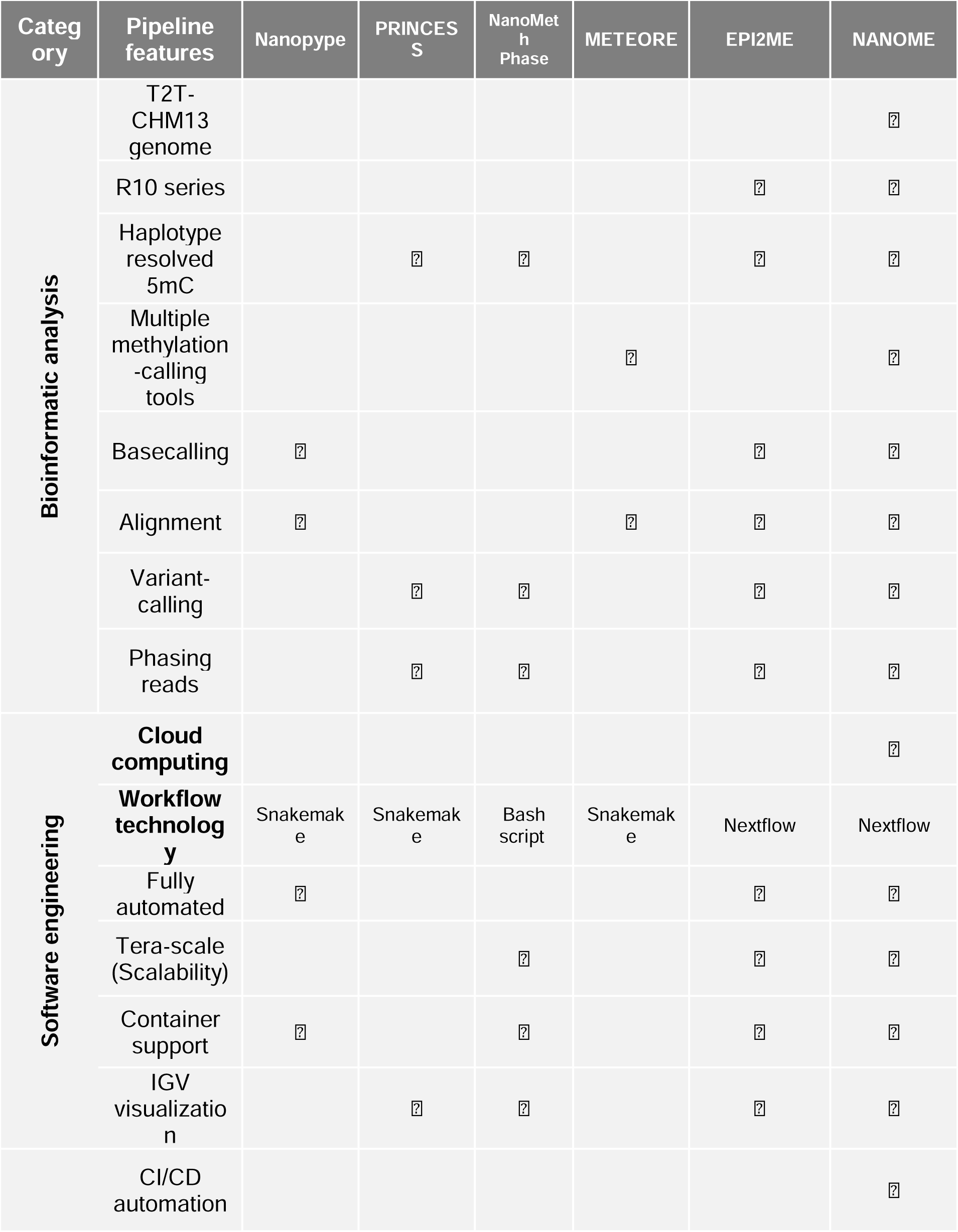
Comparison of existing nanopore sequencing DNAme pipelines, including Nanopype [35], PRINCESS [41], NanoMethPhase [42], METEORE [37], and EPI2ME(https://github.com/epi2me-labs/wf-human-variation).

### NANOME increases CpG calling accuracy via consensus pooling of state-of-the-art models

To design a methylation-calling approach using state-of-the-art tools and provide a consensus analysis of better performers, we trained an XGBoost model, which based on a distributed gradient-boosted decision tree machine learning library, on read level predictions of Nanopolish, Megalodon and DeepSignal for fully methylated and unmethylated CpG sites in chr1 of NA12878, and evaluated the model’s performance on other chromosomes (chr2-chr22, chrX and chrY) of NA12878 and APL datasets (**Figure 2A**). The input of consensus model composed of predictions of the three tools together with a 17-length sequence feature. We assigned a higher weightiness to instances observed in challenging genomic regions, i.e., discordant non-singleton DNAme pattern regions, by inverse class frequency to account for low prediction performance in these regions. The XGBoost consensus model can work on missing values, i.e., it can utilize any predictions generated by one or more tools. The consensus model outperforms Megalodon, DeepSignal and Nanopolish, with higher accuracy and coverage (**Figure 2B-D**). At discordant regions, the read level F1 score of the consensus model was 4% higher than Megalodon, DeepSignal and Nanopolish (**Figure 2B** and **Table S1**), and the site level mean square error (MSE) for the consensus model was 12% lower than these three tools (**Figure 2C** and **Table S2)**, indicating that consensus model can furtherly improve the methylation prediction performance. Interestingly, we found that the aggregating read level performance can contribute dramatically to improve site level performance by up to 11% at genome wide (**Table S2**). Additionally, the consensus model can cover 200,000 more CpGs than Nanopolish, Megalodon and DeepSignal in genome wide (**Figure 2D** and **Table S3).**

**Figure 2.**
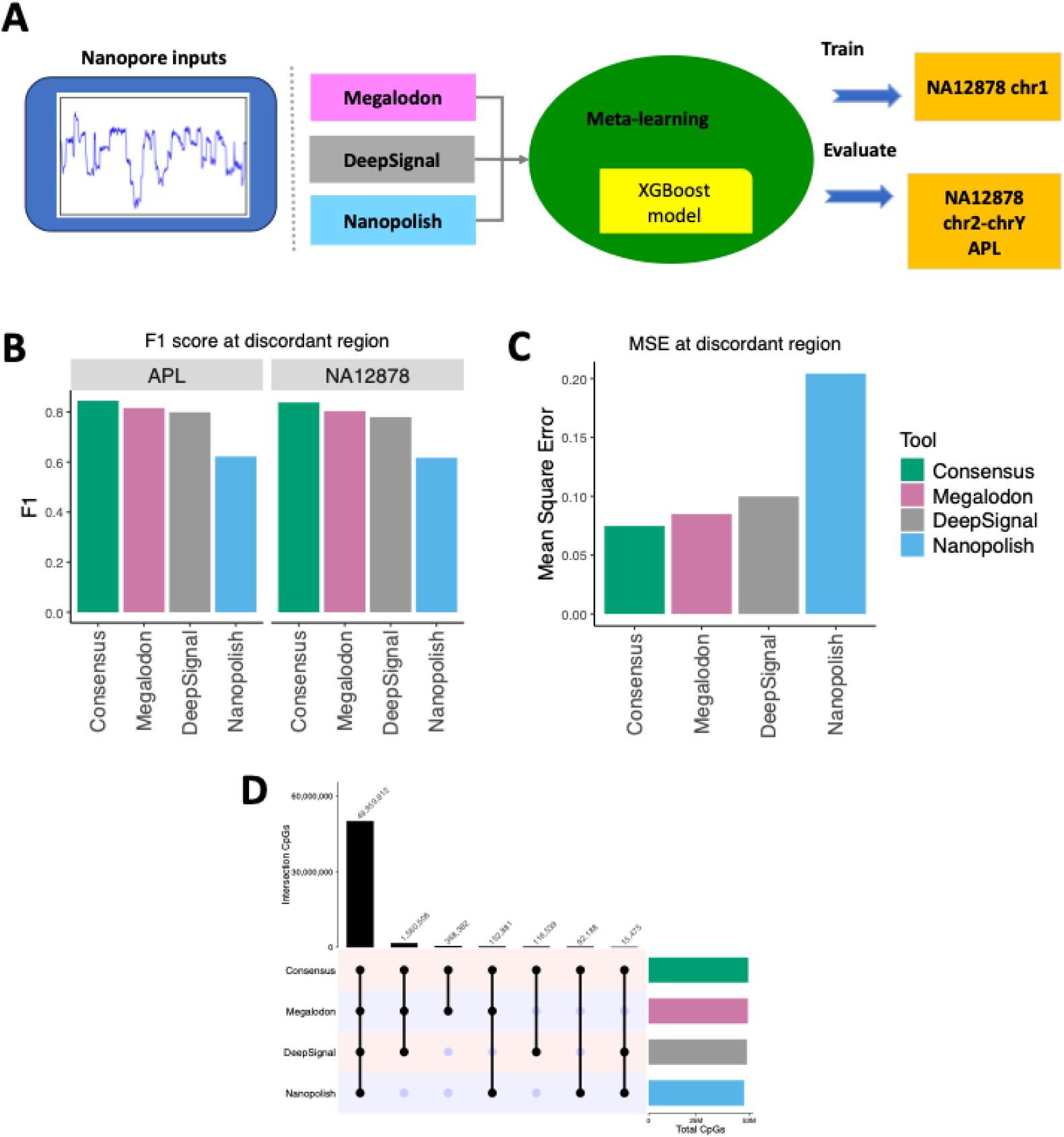
Performance comparison of consensus model on APL and chromosomes 2-22, X and Y of NA12878 data. (**A**) The Consensus model was designed in NANOME, while it uses XGBoost model trained on class weightiness assigned by inverse class frequency as weightiness with three tool’s prediction outputs and 17-length sequence feature, respectively. (**B**) Comparison of read-level performance at discordant regions on APL and NA12878. (**C**) Comparison of Mean Squared Error (MSE) for site-level discordant regions on NA12878. (**D**) CpG coverage overlapping for four tools on NA12878.

### NANOME cost-effectively optimizes coverage requirements for long-read nanopore sequencing

To optimize costs in cloud computing, we evaluated a reasonable coverage for ONT nanopore sequencing data that can achieve comparable prediction performance for consensus methylation-calling using the NA19240 and NA12878 datasets. We extracted reads at 25x coverage in using NANOME. We then down sampled the predictions for Megalodon into 5x, 10x, 15x, 20x, and 25x coverage levels. Next, we evaluated both concordance/consistency using the Pearson’s correlation coefficients (PCC) and precision through MSE at genome-wide and promoter regions between down-sampling predictions and bisulfite sequencing (BS-seq) (coverage ≥ 5). Additionally, we classified CpGs into three different methylation levels based on BS-seq methylation level: low (0, 0.33), medium (0,33, 0.67), and high (0.67, 1.0).

The genome-wide and promoter results for NA19240 and NA12878 are shown in **Figure 3A-B** and **Table S4**. We found that the PCC was higher for the promoter than the genome-wide region. When we used read coverage < 15x, concordance and precision dramatically improved compared to a read coverage of 5x for both datasets. Methylation-based sub-analyses revealed that PCC trended in the order low methylation > high methylation > medium methylation for both the NA19240 and NA12878 datasets. Overall, these findings suggest that NANOME can reduce the read coverage and maintain comparable prediction performance of ONT nanopore sequencing by using a coverage of 15x or higher, especially for CpGs with low to high methylation levels in the promoter region.

**Figure 3.**
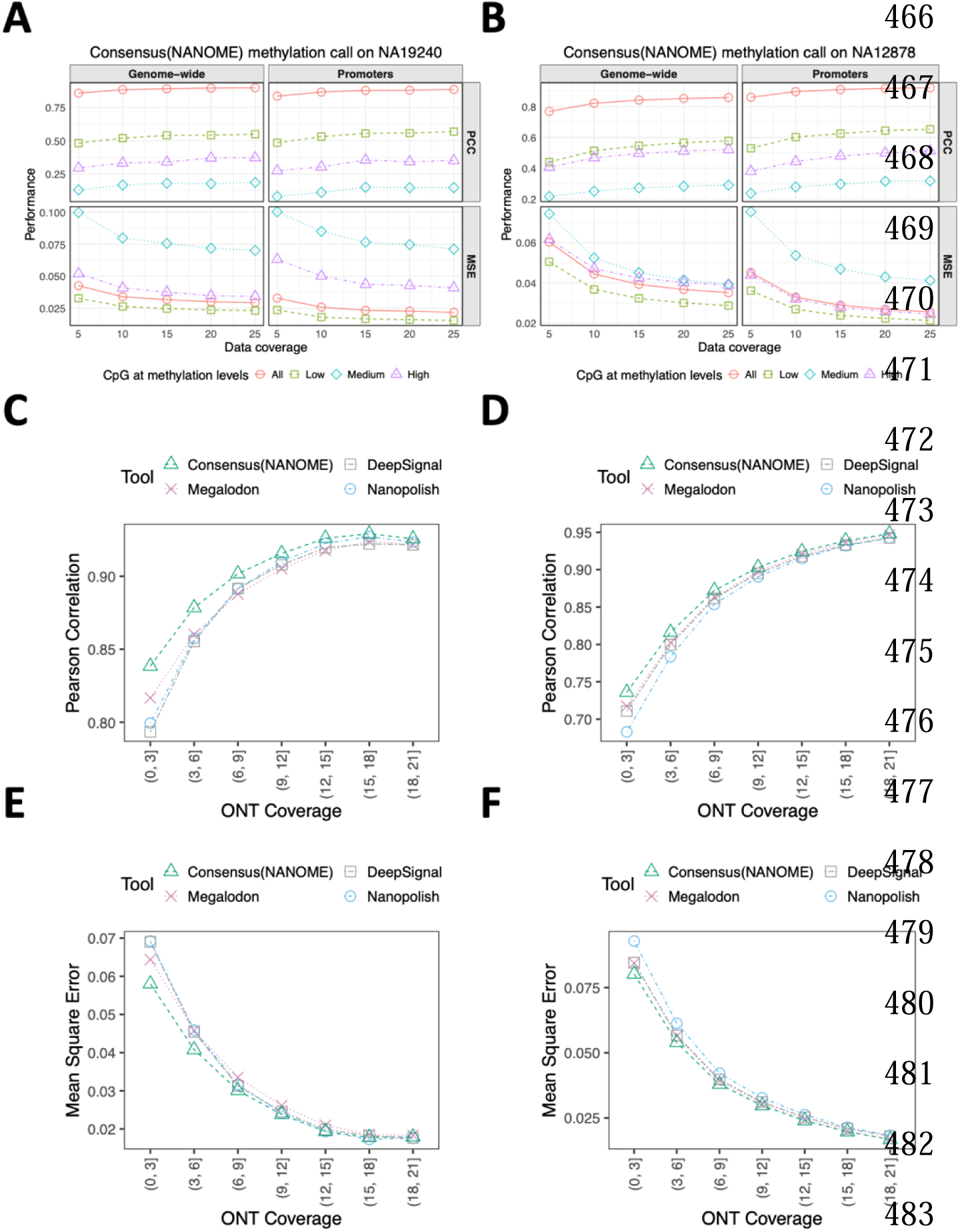
Evaluating read coverage of different basecalling tools using Pearson’s correlation coefficient (concordance) and mean square error (precision) on the NA19240 and NA12878 datasets. (**A-B**) Site-level performance using ONT’s read coverage on NA19240 (**A**) and NA12878 (**B**) for NANOME at different levels of coverage (5x, 10x, 15x, 20x and 25x). (**C-D**) Concordance comparison of the four basecalling tools at different coverage for dataset NA19240 (**C**) and NA12878 (**D**). (**E-F)** Precision comparison of the four basecalling tools at different coverages for dataset NA19240 (**E)** and NA12878 (**F**).

We further combined the predictions from four ONT methylation-calling tools and divided the ONT coverage into discrete bins—[0,3), (3,6], (6,9], …, (18,21] (**Figure S1A-B**). We then compared the PCC and MSE across all coverage bins. The results consistently show that NANOME outperforms individual tools across the different coverage levels (**Figure 3C-F and Table S5**). As such, NANOME can make ONT nanopore sequencing more accessible for biomedical studies with limited genetic material.

### Support for ONT R10 series flow cells and T2T-CHM13 genome assembly

ONT sequencing uses different generations of flow cells such as R9.4.1 and the newer R10.4, which differ in pore chemistry and sequencing accuracy. Current methylation callers (e.g., Nanopolish, Megalodon, DeepSignal) were developed for the R9.4.1 data that do not apply to R10.4 data, which require basescalling models to accurately detect methylation. So far, the tools supporting basecalling/methylation detection on data obtained using R10.4 flow cells are Guppy [18] and Dorado [48], which are readily available in the NANOME pipeline. Additionally, ONT recently introduced a new raw signal data format that supports both R9.4.1 and R10.4 data entitled POD5 to replace the older FAST5 format. Given that many methylation-calling tools were developed originally to work only with FAST5, ONT provided a POD5-to-FAST5 conversion tool (https://github.com/nanoporetech/pod5-file-format), allowing POD5-based runs to be compatible within NANOME’s pipeline. While NANOME supports the alignment and analysis using the whole human genome reference build T2T-CHM13 (v2.0) [49], offering improved mapping accuracy over older reference builds, it also supports older reference builds including HG38 (also known as GRCh38). This offers the user usage flexibility depending on the specific research question, available resources, personal preference, and community standards.

To demonstrate compatibility, we used NANOME to perform haplotype-aware methylation calling on the T2T-CHM13 genome for datasets NA12878 and NA24385 and identified regions of hypomethylation and hypermethylation (**Figure 4A-B**). We observed that the density of differentially methylated regions (DMRs) to be lower in an unresolved (gap) region compared to non-gap regions (**Figure 4C-D**). However, we observed a trend toward higher DMRs density in the unresolved regions of NA12878’s chromosomes 2, 10, and 20 (**Figure 4E**) and NA24385’s chromosome 19 (**Figure 4F**) compared to non-gap regions, but this did not quite reach significance (**Figure S2**).

**Figure 4.**
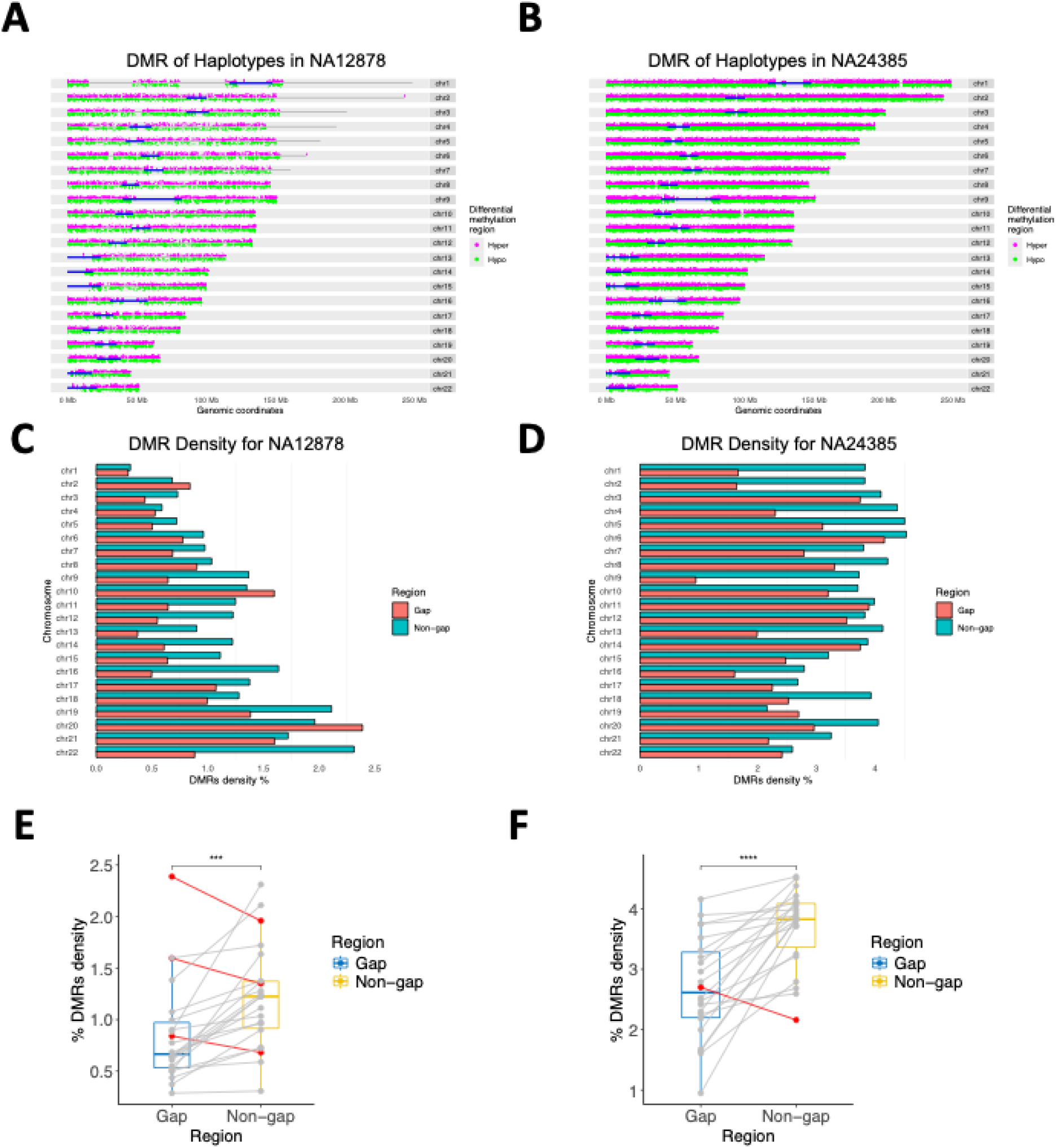
Differential methylation regions (DMRs) on T2T genome using two datasets (NA12878 and NA24385) (**A-B**) DMR density distribution for each chromosome for datasets NA12878 and NA24385. (**C-F**) DMR density distribution for NA12878 (**C, E**) and NA24385 (**D, F**) by gap state.

### Haplotype-aware allele-specific methylation for nanopore data shows improved performance

To enhance methylation phasing performance with ONT long-read nanopore data, we integrated the new variant calling module, Clair3 [50], and standardized read-level formats across tools in the NANOME pipeline. We used NA19240 to detect hayplotypes on nanopore data and compared the absolute methylation level difference with NanoMethPhase [42] between two of the detected haplotypes (**Figure 5A-B**). As shown in **Table S6**, in many imprinting control regions, the performance of methylation phasing, i.e., the absolute difference of two haplotypes by the NANOME pipeline (36.6), was comparable to the results reported by both NanoMethPhase (36.3) and Trio phasing (36.4). This is notable as Trio phasing relies on parental genotype information to resolve haplotypes, whereas NANOME achieves similar accuracy using only single sample nanopore data, making it more broadly applicable as parental genotype information is difficult to obtain.

**Figure 5.**
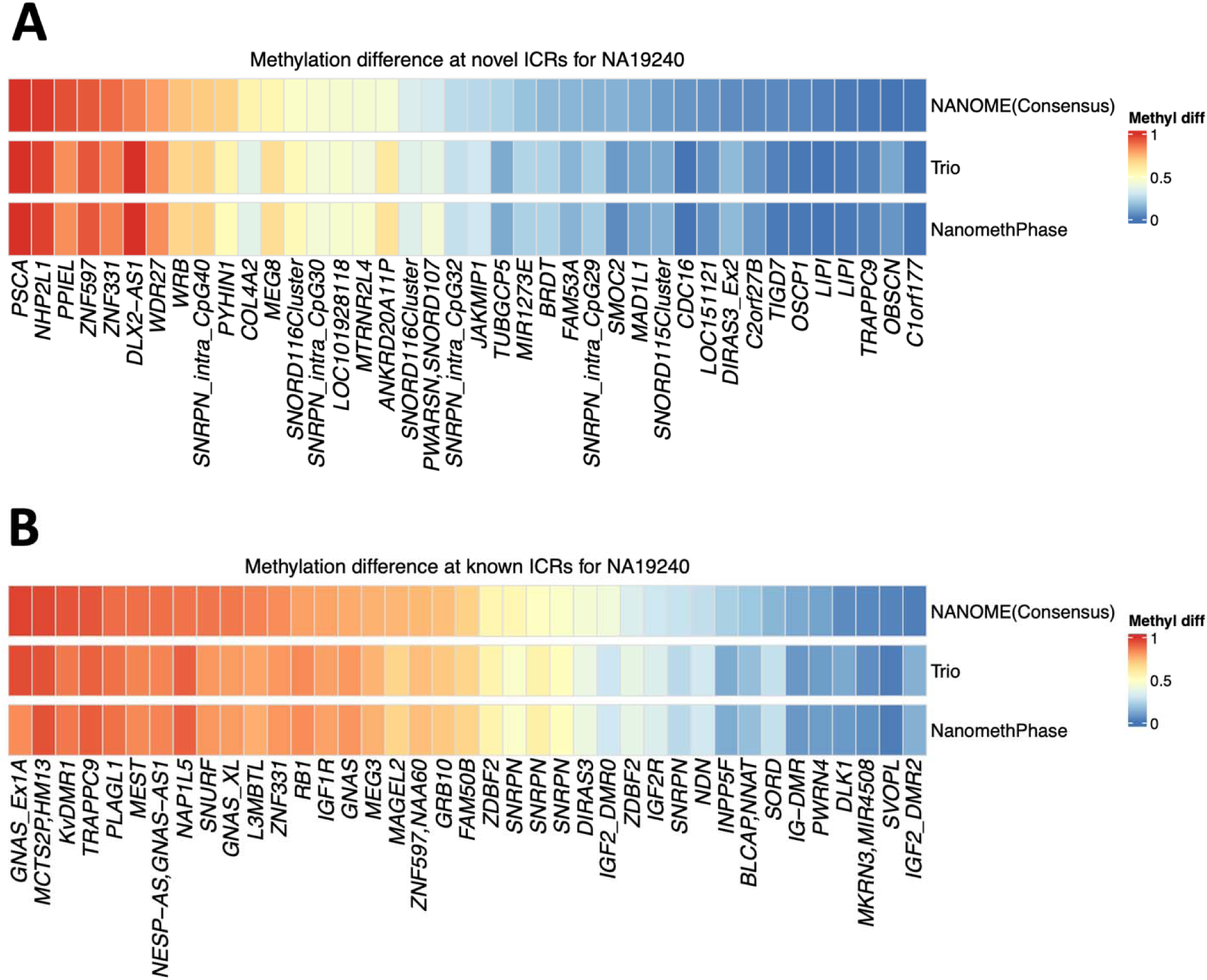
Methylation levels difference of phased CpGs at human novel Imprinting Control Regions. **(A-B)** Absolute methylation difference of HP1 and HP2 at novel (**A**) and known (**B**) imprinting control regions. We present methylation levels difference of phased CpGs, with haplotypes (HP1 and HP2) of origin at reported ICRs in the form of heatmap, similar to NanomethPhase. Trio represents the difference in methylation levels between two haplotypes for trio phasing, while NanomethPhase represents the methylation level difference for NanomethPhase.

### NANOME allows scalable and elastic computing for HPC clusters and cloud computing platforms

To address the varying hardware and software requirements of different tools, we offer Docker and Singularity support for executing our pipeline on various computing infrastructures. Additionally, we use Nextflow to provide users with the flexibility to configure the pipeline to run on both HPC clusters and the Google Cloud Computing platform. Our pipeline can parallelize any combination of tools on multiple datasets (**Figure S3A-B)**. The Nextflow implementation of NANOME simplifies the addition, management, and execution of new modules using a plain-text configuration file. We anticipate the rapid development of modules for DNA methylation-calling tools on ONT data, establishing a technical foundation for future evaluations. We do not expect that users will have to manage an overwhelming number of algorithms to identify best practices. The NANOME pipeline is built on widely accepted software standards. It integrates with any programming language through the use of software containers [51], making it possible for developers to add new modules and models without needing to modify the main pipeline codes.

## Discussion

Long-read nanopore sequencing has become a powerful tool for epigenetic analysis, particularly for detecting DNA methylation. While several pipelines for methylation detection have been published (REF), they have yet to fully automate the process, support state-of-the-art tools, incorporate phasing, and provide support for cloud platforms. We have developed NANOME, the first pipeline to offer all these key features in a single framework for ONT data analysis (**Figure 1**). While some features of other pipelines may overlap with NANOME, the reproducibility, execution platform, seamless support of online input data, and ease of use set NANOME apart. With a focus on haplotype-aware methylation detection, NANOME offers a reproducible and scalable workflow with minimal input parameters. Its implementation in the Nextflow framework allows for out-of-the-box support for various batch schedulers (SGE, PBS, and SLURM) and cloud platforms (Google Cloud, Amazon AWS), making it a versatile and accessible tool for epigenetic analysis.

NANOME is a fully automated pipeline for CpG methylation calling on ONT long-read data that integrates state-of-the-art tools (Megalodon, Nanopolish and DeepSignal) and combines their outputs through a consensus XGBoost model (**Figures 1, 2A**). It improves methylation prediction accuracy and coverage, outperforming individual tools with higher read-level F1 scores, lower site-level error, and broader CpG coverage (**Figures 2B-D; Tables S1–S3**). NANOME also enables cost-efficient sequencing by maintaining strong concordance with bisulfite sequencing at reduced coverage, particularly ≥15×, with reliable concordance and precision across coverage bins (**Figures 3; Tables S4–S5**). It supports newer nanopore formats (e.g., POD5, R10.4), allows alignment to the T2T-CHM13 genome, and detects differentially methylated regions across resolved and unresolved genomic regions (**Figures 5, S2**). By integrating Clair3, NANOME enables haplotype-aware methylation analysis using only single-sample data with performance comparable to trio-based approaches (**Figure 5; Table S6**). Built on Nextflow, NANOME supports cloud and HPC deployment, offering scalable, parallel, and modular analysis workflows across platforms (**Figure S3**).

Telomere shortening, a process that is accelerated in some diseases and lifestyle conditions, is a well-established biomarker of aging [52]. Our pipeline is not only capable of supporting state-of-the-art tools but also the latest T2T genomes [53]. This allows researchers to detect allele-specific methylation in even the most challenging regions of the genome. With this feature, our pipeline can improve the accuracy and precision of DNA methylation analysis, leading to a deeper understanding of epigenetic regulation in various biological processes. Therefore, NANOME significantly advances allele-specific methylation detection for long-read nanopore sequencing. The ability to support updated genomes will further enhance the utility and relevance of our pipeline in the rapidly evolving field of epigenetic research.

The development of long-read nanopore 5mC methylation detection methods have significantly advanced our understanding of the biological role of 5mC. These methods have provided researchers with the ability to directly sequence and detect 5mC modifications, which has shed new light on the importance of this epigenetic mark in various biological processes [3]. With these technologies, scientists can now investigate the role of 5mC in gene regulation, cellular differentiation, and disease states. The ability to detect and analyze 5mC modifications with high accuracy and sensitivity has paved the way for discoveries in epigenetic research and may ultimately lead to novel therapeutic approaches for various diseases. In addition, recent advances in nanopore sequencing technology have led to the development of various methods for predicting DNA methylation. Among these methods, consensus prediction using individual reads has emerged as a promising approach. The key to the success of the consensus model lies in its meta-learning mechanism, which is based on top performers, and the feature of training data that contain both concordant and discordant CpG sites. By leveraging these features, the model can effectively learn from the strengths of different methods and improve its prediction accuracy.

We are working toward having a full-day turnaround of our complete computational and cloud-compatible pipeline. By implementing real-time basecalling, methylation-calling, variant-calling, and phasing for ONT data, we could achieve de novo haplotype-aware methylation detection from DNA in one command line in 48h. This user interface level and speed would facilitate screening for epigenetic abnormalities in challenging-to-sequence regions of human genomes.

## Conclusion

NANOME is compatible with seven of the primary DNA methylation-calling tools in use today and is specifically designed to provide more robust results based on the top three performers. Some tools included in NANOME, such as Megalodon, DeepSignal, and Guppy, rely on GPU-based software. The accuracy and coverage of CpG sites of NANOME consensus algorithms are dependent on read-level predictions from the top three performers (Nanopolish, Megalodon, and DeepSignal), which can provide more accurate results. Looking ahead, we anticipate the development of modules focused on four primary tasks: (1) basecalling, (2) quality control, (3) alignment, and (4) methylation calling. However, we do not expect users to manage all four processes continuously. NANOME provides a technical foundation for such evaluations, and its adoption can enable community-wide development of DNA methylation-calling for analyzing ONT nanopore sequencing data. The pipeline is based on widely accepted software standards and interoperates with any programming language through software containers, enabling developers to add new modules and models. Our experience suggests that new users can master the Nextflow command line interface or the Web UI within a single training day.

## Methods

### Oxford Nanopore sequencing data

ONT sequencing data for NA12878 is available as an Amazon Web Services Open Data Set at https://github.com/nanopore-wgs-consortium/NA12878 [54]. ONT sequencing data for NA19240 is available at ENA (https://www.ebi.ac.uk/ena/browser/home) under the accession number PRJEB26791 [55]. ONT sequencing data for NA24385 is available from the Oxford Nanopore Technologies Open Data repository, hosted on Amazon Web Services (AWS) S3, at the following location: s3://ont-open-data/gm24385_q20_2021.10/.

All three datasets are sequenced using high-throughput-whole genome sequencing from DNA extracted from Epstein-Barr virus-transformed B-lymphocyte cell lines, established by the Coriell Institute as part of the International HapMap and 1000 Genome Projects. NA19240 comes from a female with African ancestry (age NA), NA12878 comes from a female with European ancestry (age NA) and NA24385 comes from a male with European ancestry (age = 45 at sampling). ONT sequencing data for APL is available at the European Genome-phenome Archive (EGA) under accession number EGAS00001005613 [38].

### DNA methylation data and preprocessing

Whole genome bisulfite sequencing (WGBS) and reduced representation bisulfite sequencing (RRBS) files are downloaded from the ENCODE portal (https://www.encodeproject.org) with the following identifiers: ENCFF835NTC and ENCFF279HCL (WGBS for NA12878), ENCFF000LZS and ENCFF000LZT (RRBS for NA19240). WGBS data for APL is available at the EGA under accession number EGAS00001005610. For RRBS data, adapters were trimmed using TrimGalore (http://www.bioinformatics.babraham.ac.uk/projects/trim_galore/) followed by base quality checking using FastQC (http://www.bioinformatics.babraham.ac.uk/projects/fastqc/). Reads were then aligned to the human genome reference hg38 (GCA_000001405.15) using Bismark before removing duplicate reads. To determine the DNA methylation status at each CpG site, we ran Bismark scripts (https://github.com/TheJacksonLaboratory/BS-seq-pipleine) and obtained the union of CpG sites from two replicates (WGBS or RRBS) as the corresponding DNA methylation ground truth. We use CpGs with coverage≥5 in WGBS or RRBS for performance evaluation.

### Basecalling and methylation-calling for ONT nanopore sequencing

The NANOME pipeline offers support for achieving consensus using three different tools, namely Nanopolish (v0.13.2), Megalodon (v2.4.2), and DeepSignal (v0.1.10). For APL, NA12878 and NA19240 R9.4.1 data, basecall was using Guppy model dna_r9.4.1_450bps_hac.cfg. We then performed methylation calling for above three tools and the NANOME consensus model. These tools utilize different algorithms and are trained to detect DNA methylation modifications in distinct ways. Nanopolish uses a Hidden Markov Model (HMM) to call 5mCs in a CpG context, assigning a log-likelihood ratio (LLR) for each CpG site. It detects methylated CpGs (LLR > 0) and unmethylated CpGs (LLR < 0) at the read level. Megalodon is a novel command-line tool developed by ONT that leverages a recurrent neural network (RNN) model based on deep learning to identify modified bases. DeepSignal uses a Bidirectional Long/Short-Term Memory (BiLSTM) and Inception structure to detect the methylation state. For NA24385 R10.4 data, NANOME pipeline performs the methylation calling using Guppy with R10.4 basecall and methylation calling models: dna_r10.4_e8.1_hac.cfg and dna_r10.4_e8.1_modbases_5mc_cg_hac.cfg. The performances of these methods, which rely on prior knowledge of expected signal deviations, are highly dependent on the training data, which typically comprises fully unmodified and fully modified samples.

### Training and testing data design

Our study utilized the fully methylated (100%) and unmethylated (0%) CpG sites with coverage≥5 from the NA12878 BS-seq dataset to train and validate our models. First, XGBoost was applied to chromosome 1 of the NA12878 data for 5mC classification using 3-fold cross-validation. All methods were then tested on other chromosomes of NA12878 and APL compared with WGBS. For read-level evaluation, we joined all CpG predictions for each read from each tool and compared their accuracy, F1-score, precision, recall, and ROC-AUC on fully methylated or unmethylated CpG sites for chr2-chr22, chrX, and chrY of both NA12878 WGBS and APL WGBS (**Table S1**). Additionally, for site-level evaluation, we aggregated CpG methylation frequency using joint predictions, i.e., read-level predictions generated by all tools, and compared the Pearson correlation coefficient (PCC) and mean square error (MSE) between predicted frequency and methylation level in BS-seq for all CpG sites with coverage≥5 in the NA12878 WGBS dataset (**Table S2**).

### XGBoost hyper-parameter selection for consensus model and 5mC prediction

To build our XGBoost classifier, we utilized the XGBoost Python library and conducted experiments with various hyperparameter settings for the XGBoost model. We performed hyperparameter selection on key parameters such as the learning rate, number of estimators, maximum tree depth, subsample ratio, and L1 and L2 regularization terms on weights. Then, we fine-tuned the network hyperparameters through a random search to obtain the best possible values. The hyperparameter settings and their respective experimental values are listed below, with the selected values indicated in bold. Our final selected values achieved an F1-score of 0.926:

i. learning_rate: 0.01, 0.05, **0.1** and 0.2
ii. n_estimators: 5, 10, 20, 40 and **60**
iii. max_depth: 3, 6 and **9**
iv. subsample: 0.6, **0.8** and 1
v. colsample_bytree: 0.6, 0.8 and **1**
vi. reg_alpha: **0**, 0.5, 2 and 5
vii. reg_lambda: **0.5**, 1 and 2

### Differential methylation regions

To identify DMRs across phased two haplotypes, we employed the methylKit package (https://github.com/al2na/methylKit), a comprehensive R tool designed for the analysis of methylation data [56]. The analysis was structured around the utilization of a sliding window approach, with both the window size and step size specifically configured to 1,000 base pairs. DMRs were subsequently delineated based on stringent criteria: only regions exhibiting a methylation difference exceeding 25% were considered, ensuring that only substantial alterations in methylation patterns were flagged. Additionally, to control for the false discovery rate inherent in multiple testing, a threshold was set wherein only regions with a Chi-squared test q-value below 0.05 were classified as differentially methylated. This dual-threshold approach integrates both biological significance and statistical rigor, facilitating the reliable identification of regions where methylation status is associated with haplotype phase.

## Code Availability

The NANOME pipeline code is available in GitHub: https://github.com/LabShengLi/nanome.

## Acknowledgements

Dr. Sheng Li is supported by the following grants: R35GM133562, U01HG013175, U01CA271830. We would also like to express our sincere gratitude to scientific writer Anna Lisa Lucido from The Jackson Laboratory, whose invaluable contributions significantly enhanced the quality of this manuscript.

## Supplementary Files

Supplementary Figures S1-S3.

Supplementary Tables S1-S6.

